# Stability of subsystem solutions in agent-based models

**DOI:** 10.1101/191106

**Authors:** Matjaž Perc

**Author notes:** Electronic address, Home page: www.matjazperc.com.

## Abstract

The fact that relatively simple entities, such as particles or neurons, or even ants or bees or humans, give rise to fascinatingly complex behavior when interacting in large numbers is the hallmark of complex systems science. Agent-based models are frequently employed for modeling and obtaining a predictive understanding of complex systems. Since the sheer number of equations that describe the behavior of an entire agent-based model often makes it impossible to solve such models exactly, Monte Carlo simulation methods must be used for the analysis. However, unlike pairwise interactions among particles that typically govern solid-state physics systems, interactions among agents that describe systems in biology, sociology or the humanities often involve group interactions, and they also involve a larger number of possible states even for the most simplified description of reality. This begets the question: When can we be certain that an observed simulation outcome of an agent-based model is actually stable and valid in the large system-size limit? The latter is key for the correct determination of phase transitions between different stable solutions, and for the understanding of the underlying microscopic processes that led to these phase transitions. We show that a satisfactory answer can only be obtained by means of a complete stability analysis of subsystem solutions. A subsystem solution can be formed by any subset of all possible agent states. The winner between two subsystem solutions can be determined by the average moving direction of the invasion front that separates them, yet it is crucial that the competing subsystem solutions are characterized by a proper composition and spatiotemporal structure before the competition starts. We use the spatial public goods game with diverse tolerance as an example, but the approach has relevance for a wide variety of agent-based models.

## I. INTRODUCTION

The total is more than just the sum of all the parts. This old adage applies as well to anthills as it does to neurons that form our brain, and in simple terms it describes the essence of complex systems. The often beautiful and fascinating collective behavior that emerges as a result of interactions between a large number of relatively simple entities has been brought to the point by the physicist and Nobel laureate Philipp Anderson in his paper *More is different* [1], which is often cited as the birthstone of complex systems science, at least for physicists. An important point is that the collective behavior entails emergent phenomena that can hardly, if at all, be inferred from the properties of the individual parts [2, 3]. In the decades past, however, the words complex and complexity have been used to describe all kinds of systems and phenomena, within and outside of physics, from computer science to politics, to the point today where what exactly is complex systems science is anything but easy to put shortly. Thankfully, a recent review by Yurij Holovatch, Ralph Kenna and Stefan Thurner titled *Complex systems: Physics beyond physics* [4] reaffirms what ought to be the essence of complex systems from a physicist point of view, and it clarifies what makes them conceptually different from systems that are traditionally studied in physics. Further to that effect, we have the *Focus on Complexity* collection to be published in the *European Journal of Physics*, to which this paper belongs.

Regardless of the definitions, it is thoroughly established that methods of statistical physics, in particular Monte Carlo methods and the theory of collective behavior of interacting particles near phase transition points – a classical subject that is thoroughly covered in comprehensive reviews [5–7] and books [8–11] – have proven to be very valuable for the study of complex systems. In fact, these methods have been applied to subjects that, in the traditional sense, could be considered as out of scope of physics. Statistical physics of social dynamics [12, 13], of human cooperation [14, 15], of spatial evolutionary games [16–21], of crime [22], and of epidemic processes and vaccination [23, 24], are all examples of this exciting development. The advent of network science as an independent research field can also be considered as an integral part of this development [25–33], providing models and methods that have revived not just statistical physics, but also helped complex systems science to grow.

Agent-based models constitute a red thread through much of complex systems research. The obvious idea is that agents interact with one another, typically through an interaction network. And while agents themselves are simple entities that can typically choose only between a couple of different states, their interactions give rise to collective behavior that could not possibly be anticipated from the simplicity of each individual agent. Agent-based modeling has been used for simulating and studying human and social systems [34, 35], ecological systems [36], economic systems [37, 38], as well as evolution and cooperation [39, 40], to name just some examples. There are reviews [4] and tutorials [41] available, which cover the subject in much detail.

We are here concerned with a much too frequently overlooked but, especially from the viewpoint of physics, very important aspect of agent-based modeling, namely the validity and stability of observed simulation outcomes. In principle, any agent-based model is fairly easy to simulate with a Monte Carlo simulation method. Realistically, however, the acquisition of correct results requires a careful approach that is seldomly used and advocated for. The root of the problem lies in the fact that any subsystem solution can be the solution of the whole system. Subsystem solutions are simply solutions that are formed by a subset of all possible agent states. Evidently, if an individual agent can choose between three or more different states (not just two, like spin up/down), the number of possible subsystem solutions increases fast, and therewith also the severity of the problem. To determine the stability of subsystem solutions, and thus to determine the most stable system-wide solution, we must perform a systematic stability check between all unique pairs of subsystem solutions, as determined by the average moving direction of the invasion front that separates them. Only then can we be certain that the simulation outcome of an agent-based model is actually stable and valid in the large system-size limit, and we can proceed with the determination of phase transitions that separate different stable solutions, and with the determination of the responsible microscopic processes. Since this approach has relevance for a wide variety of agent-based models – in fact for all agent-based models where there are more than two possible agent states or competing strategies – the methodology is of the utmost important for those that teach as well as for those that learn and practice this aspect of complex systems research.

In what follows, we first introduce the spatial public goods game with diverse tolerance as the example agent-based model in Section II, which we will then use to didactically demonstrate the stability of subsystem solutions in Section III. Lastly, we sum up and discuss the relevance of the described approach for agent-based models in different fields of research in Section IV.

## II. PUBLIC GOODS GAME WITH DIVERSE TOLERANCE

The public goods game is simple and intuitive [42], and it is in fact a generalization of the pairwise prisoner’s dilemma game to group interactions [43]. In a group of agents, each one can decide whether to cooperate or defect. Cooperators contribute *c* =1 to the common pool, while defectors contribute nothing. The sum of all contributions is multiplied by a multiplication factor *r* > 1, which takes into account synergistic effects of cooperation. In particular, there is an added value to a joint effort that is often more than just the sum of individual contributions. After the multiplication, the resulting amount of public goods is divided equally amongst all group members, irrespective of their strategy. In a group *g* containing *G* agents the resulting payoffs are thus

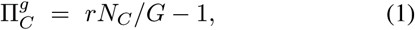

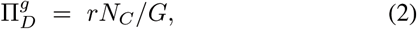

where *N*_*C*_ is the number of cooperators in the group. Evidently, the payoff of a defector is always larger than the payoff of a cooperator, if only *r* < *G*. With a single parameter, the public goods game hence captures the essence of a social dilemma in that defection yields highest short-term individual payoffs, while cooperation is optimal for the society as a whole.

To add tolerance to this basic setup, we have, in addition to cooperators (*C*) and defectors (*D*), also loners (*L*) and tolerant agents (*M*_*i*_) [44,45]. Loners are agents that simply abstain from the game and settle for a small but secure payoff σ = 1. Tolerant agents, on the other hand, either cooperate or abstain from the game depending on the number of defectors *i* within a group. There are as many levels of tolerance as there are possible defectors in the group, so that *i* = 0,…, *G* – 1. If the number of defectors in a group is smaller than *i* the agent *M*_*i*_ acts as a cooperator. Otherwise it acts as a loner. Accordingly, the higher the value of *i*, the higher the number of defectors that are tolerated by the corresponding agent within a group. As the two extreme cases, *i* = 0 implies that the agent will always act as a loner, while *i* = *G* – 1 indicates that the agent will act as a loner only if all other group members are defectors. Importantly, regardless of the choice an *M*_*i*_ agent makes, it always bears the cost *γ* > 0 as a compensation for knowing the number of defectors in a group, i.e., the cost of inspection. Also importantly, the *r* > 1 factor is applied only if there are at least two contributions made to the common pool from within the group. Otherwise, a lonely contributor is unable to utilize on the synergistic effect of a group effort, and hence *r* = 1 applies.

For the mathematical formulation of the payoffs, it is convenient to introduce *δ*_*i*_ = 0 if *N*_*D*_ ≥ *i* and *δ*_*i*_ = 1 if *N*_*D*_ < *i*, where *N*_*D*_ is the number of defectors in a group. The total number of contributors to the common pool then becomes

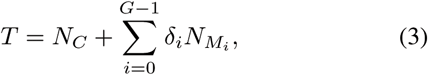

where *N*_*s*_ denotes the number of agents in the group who follow strategy s. By using this notation, the payoff values of the competing strategies obtained from each group *g* are

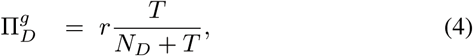

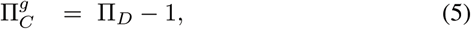

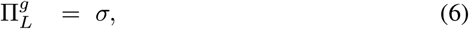

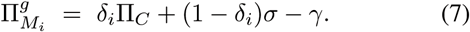

The public goods game is here staged on a square lattice with periodic boundary conditions where *L*^2^ agents are arranged into overlapping groups of size *G* = 5 such that everyone is connected to its *G* – 1 nearest neighbors [42]. Accordingly, each individual belongs to *g* = 1, . . . , *G* different groups. With these choices, we thus have an agent-based model where each agent can choose between *n* = 8 different states/strategies, namely *C*, *D*, *L*, *M*_0_, *M*_1_, *M*_2_, *M*_3_ and *M*_4_. In order to determine the number of all possible subsystem solutions, we have to determine the number of combinations without repetition when the number of strategies in a subsystem can be any between 1 ≤ *k* ≤ 8. We thus have a total of 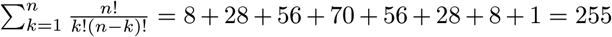 possible subsystem solutions, and no less than 32385 unique pairs to compete against each other in a round-robin tournament. Thankfully, apart from single-state subsystem solutions, which are all trivially stable, the large majority of other *k*–state subsystem solutions, where *k* > 1, turn out to be unstable in the *r* – *γ* parameter plane. This significantly reduces the effort that is needed to determine the most stable systemwide solutions and the phase transitions that separate them.

Monte Carlo simulations are carried out comprising the following elementary steps. A randomly selected agent *x* with strategy *s*_*x*_ plays the public goods game in all the *g* = 1,…, *G* groups where it is member. Its overall payoff Π_*s*_*x*__ is the sum of all the payoffs 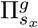 acquired in each individual group. Next, one randomly chosen neighbor of agent *x* also acquires its payoff *Π*_*s*_*y*__ in the same way. Lastly, player *y* adopts the strategy from player *x* with a probability given by the Fermi function

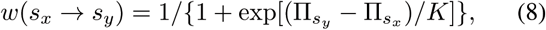

where *K* = 0.5 quantifies the uncertainty by strategy adoptions [46], implying that the strategies of agents with higher payoffs are readily adopted, although the opposite is not impossible either. In agreement with the random sequential update, each full Monte Carlo step (MCS) gives a chance to every player to change its strategy once on average. The average fraction of each state/strategy in the system (*ρ*_*s*_*x*__) is determined in the stationary state after a sufficiently long relaxation time is discarded, that is when the average fraction of the strategies becomes time independent.

## III. RESULTS

Before turning to the stability of subsystem solutions, a note concerning initial conditions is in order. It is often written that strategies are initially distributed uniformly at random over a lattice to give each of them the same chance of evolutionary success. Evidently, this has to do with the fact that random initial conditions make certain that each strategy occupies about the same amount of space in a population. But this alone does not confer equal chances of survival to all strategies, except if the competing strategies are only two. If the competing strategies are more than two, it is actually quite impossible to engineer ‘fair’ initial conditions because stable subsystem solutions are not made up solely of single strategies, but could also be formed by two-strategy, three-strategy, four-strategy, and so on, combinations. Importantly, these subsystem solutions first have to form, i.e., attain their actual spatiotemporal structure (for example traveling or target waves, checkerboard patterns, compact clusters, etc.) before they would begin competing against each other. But the formation of different subsystem solutions is in general characterized by different, and sometimes very different, time scales. So what random initial conditions, if paired with a very large system size, actually do accomplish, is that they give a chance to each subsystem solution to emerge somewhere locally in the population. If using small system sizes, however, only those subsystem solutions can evolve whose characteristic formation times are sufficiently short.

Since we have no way of knowing which initial configuration of strategies will yield a stable subsystem solution, our best option is thus to use random initial conditions with a very large system size, and hope for all of them to emerge at some point in time. After we identify them, however, it is much more efficient and fair in terms of equal survival chances to use prepared initial states, and to do a proper stability analysis of subsystem solutions, as we describe next on an example.

We begin in reversed order, showing first the phase diagram of the public goods game in Fig. 1, which would normally be the final result of a proper subsystem stability analysis. Different generally valid observation can be made. In the first place, it can be observed that the higher the value of *r*, the higher the tolerance can be, and vice versa. Secondly, if the cost of inspection is too high, or if the value of the synergy factor is either very low or very high, tolerant players cannot survive. Thirdly, strategies *M*_0_ and *M*_4_ never survive, indicating that fully intolerant or overly tolerant strategies are not evolutionary viable. Several more precise observations can be made, but these are presently outside of scope. For details we refer to [45], where the public goods game with diverse tolerance levels was presented and studied first.

**FIG. 1:**
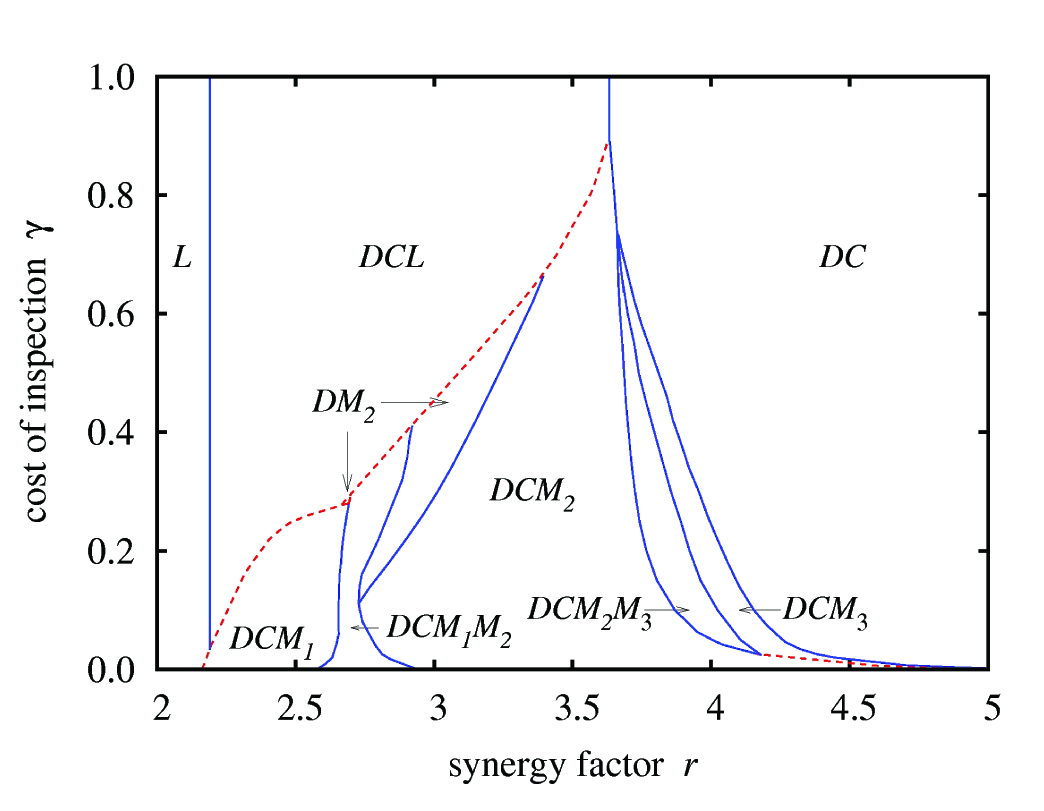
Phase diagram of the 8-strategy agent-based model on the *r* – *γ* parameter plane. Red dashed (blue solid) lines denote discontinuous (continuous) phase transitions. Of particular interest is the *DCL* → *DCM*_1_*M*_2_ discontinuous phase transition, which we will focus on in Figs. 2 and 3.

For a concrete example of the stability analysis of two subsystem solutions, we pick the *DCL* → *DCM*_1_*M*_2_ discontinuous phase transition and show how the precise value of r at which the transition occurs is determined. In general, the procedure is particularly useful (and often needed) in the vicinity of discontinuous phase transition, which can otherwise be quickly determined wrongly as a consequence of random extinctions that are due to the usage of too small system sizes. Continuous phase transitions are in this regard somewhat less demanding.

The analysis begins by first partitioning the square lattice in half, such that strategy changes across the vertical border are prohibited. Instead, periodic boundary conditions are applied from the middle of the lattice towards the left and right edge, respectively. More precisely, each separated part of the lattice (each half) has its own periodic boundary conditions. There are thus no interactions with the players from the other half, not in terms of the determination of payoffs, and also not in terms of strategy transfer. Only when the vertical border is removed (see below) do the traditional periodic boundary conditions across the whole lattice apply. Each half of the lattice is populated uniformly at random only with the strategies that form the subsystem solution of which stability we wish to determine. This is demonstrated in the leftmost panel of Fig. 2, where in the left half of the lattice we have agents *D*, *C* and *L*, while in the right half we have agents *D*, *C*, *M*_1_ and *M*_2_, randomly distributed in both cases. After a sufficiently long transient, which depends on their formation time scales, the two subsystem solutions attain their actual spatiotemporal structure, as denoted by *DCL* and *DCM*_1_*M*_2_ subscripts on top of the middle panel of Fig. 2. In the three-strategy phase strategies *D*, *C* and *L* dominate each other cyclically. The existence of the *D* → *C* → *L* → *D* closed loop of dominance is also evident from the spatiotemporal structure that manifests itself as traveling spiral-like structures in the left half of the middle panel of Fig. 2. Conversely, the four-strategy *DCM*_1_*M*_2_ phase depicted in the right half of the middle panel of Fig. 2 is somewhat more static. In fact, there typical compact clusters of cooperative strategies (especially of *C* and *M*_1_) that are exploited by more or less isolated defectors can be observed.

**FIG. 2:**
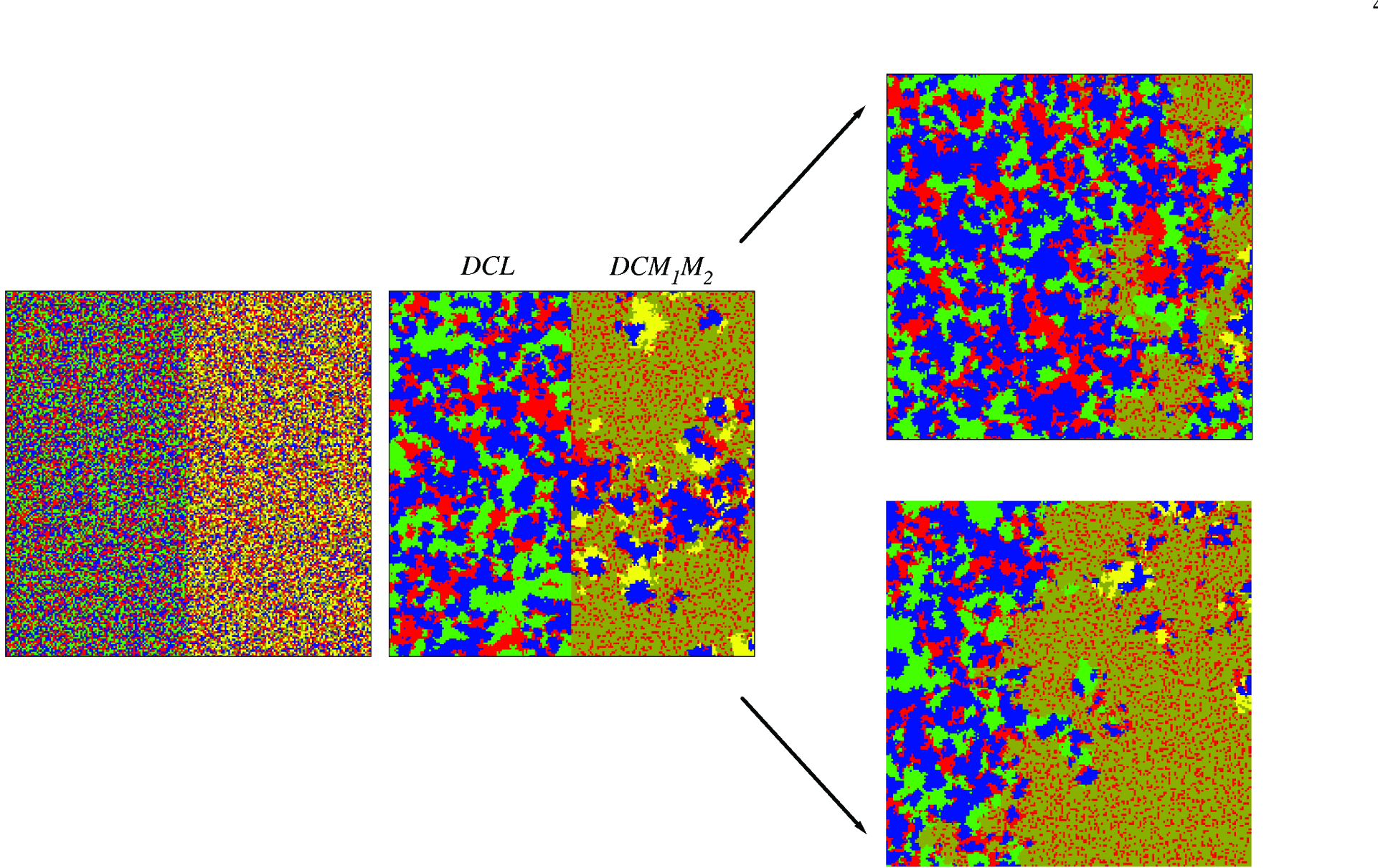
Subsystem stably analysis of two individually stable solutions, namely the three-strategy *DCL* (red, blue and green agents) and the four-strategy *DCM*_1_*M*_2_ (red, blue, yellow and dark yellow agents) phase, which are separated by a discontinuous phase transition in Fig. 1. The leftmost panel shows the initial setup, where the square lattice is vertically partitioned, such that strategy changes across the border are not allowed. In the left half we have agents *D*, *C* and *L*, and in the right half we have agents *D*, *C*, *M*_1_ and *M*_2_ – in both cases distributed uniformly at random. After 20000 MCS the two subsystem solutions attain their actual spatiotemporal structure, as denoted by *DCL* and *DCM*_1_*M*_2_ subscripts on top of the middle panel. From here on the vertical border can be removed to determine the average moving direction of the invasion front that separates the two subsystem solutions. If we use *r* = 2:80 and *γ* = 0:35, we obtain the upper right snapshot, where the DCL phase will turn out to be the winner. Conversely, if we use *r* = 2:81 and *γ* = 0:35, we obtain the lower right snapshot, where the *DCM*_1_*M*_2_ phase will turn out to be the winner. Importantly, for both values of *r* the two formed subsystem solutions in the middle panel are qualitatively exactly the same, i.e., both phases are individually stable regardless of which value of *r* is used.

After the subsystem solutions are formed, the vertical border can be removed to determine the average moving direction of the invasion front that separates the two subsystem solutions, i.e., to effectively determine the winner of the competition and thus the most stable system-wide solution. In our example depicted in Fig. 2, we obtain the upper right snapshot for *r* = 2.80 and *γ* = 0.35, where the DCL phase will turn out to be the winner, while for *r* = 2.81 and *γ* = 0.35 we obtain the lower right snapshot, where the *DCM*_1_*M*_2_ phase will turn out victorious. Importantly, for both values of *r* the two formed subsystem solutions in the middle panel are qualitatively exactly the same, and can thus be used as the starting point of the competition in both cases. In other words, both the *DCL* and the *DCM*_1_*M*_2_ subsystem solutions are individually stable regardless of which value of *r* is used. The performed subsystem stability analysis reveals, however, that for *r* = 2.80 the most stable system-wide solution is the *DCL* phase, while for *r* = 2.81 the most stable system-wide solution is the *DCM*_1_*M*_2_ phase. This also exactly determines the critical value of *r* at which the discontinuous phase transition occurs for the applied value of *γ* = 0.35. Evidently, to obtain the complete phase diagram shown in Fig. 1, this procedure should be applied in the vicinity of all phase transition points for a sufficiency fine mesh of values across the *r*–*γ* parameter plane.

We emphasize that finite-size effects can easily play an obstructive role in the described stability analysis of subsystem solutions. If we start the evolution from a random initial state using a small system size, it can easily happen that we observe a misleading evolutionary outcome, simply because the phase that would be a stable solution at a large system size has no chance to emerge – for example, one of the strategies that would be necessary to form it dies out beforehand due to the small system size. But that is not the only caveat. Even if we use prepared initial states for the stability analysis as depicted in the leftmost panel of Fig. 2, we should be careful because the part of the lattice allocated to each subsystem solution should be large enough for the latter to actually form. In fact, the fluctuations of strategies in the cyclically dominated *DCL* phase could be extremely large, which is generally valid for agent-based models that are governed by cyclic dominance, especially close to phase transitions [47]. Therefore this subsystem solution alone requires a large system size to avoid an accidental extinction of a strategy, upon which the closed loop of dominance would be broken and the subsystem solution could of course no longer emerge.

It is lastly instructive to corroborate the presented subsystem stability analysis with time courses of strategy fractions that correspond to the snapshots presented in Fig. 2. To that effect, we show in the upper panel of Fig. 3 how the fractions of strategies C, *D*, *L*, *M*_1_ and *M*_2_ change over time when using *r* = 2.80 and *γ* = 0.35. As reported above, for these parameter values the wining subsystem solution is the *DCL* phase that is governed by cyclic dominance. It can be observed that, as soon as the vertical border at 20000 MCS is removed (arrow), the fractions of strategies *M*_1_ and *M*_2_ start declining. Evidently, the invasion front that separates the two subsystem solutions is moving into the *DCM*_1_*M*_2_ phase, thus indicating that the *DCL* will eventually win. Indeed, at around 50000 MCS the two tolerant strategies die out. Conversely, the lower panel of Fig. 3 shows how the fractions of the five strategies change over time when using *r* = 2.81 and *γ* = 0.35. Here at 20000 MCS (arrow) the fraction of strategy *L* starts declining, until it eventually vanishes completely at around 55000 MCS. The fraction of strategy *C* also starts declining when the border is removed, but as soon as the strategy L dies out, it saturates to a non-zero value, thus giving rise to the victory of the *DCM*_1_*M*_2_ phase. Results presented in Fig. 3 thus demonstrate how the time courses reveal the average moving direction of the invasion front, and thus help determine the winner between two subsystem solutions.

**FIG. 3:**
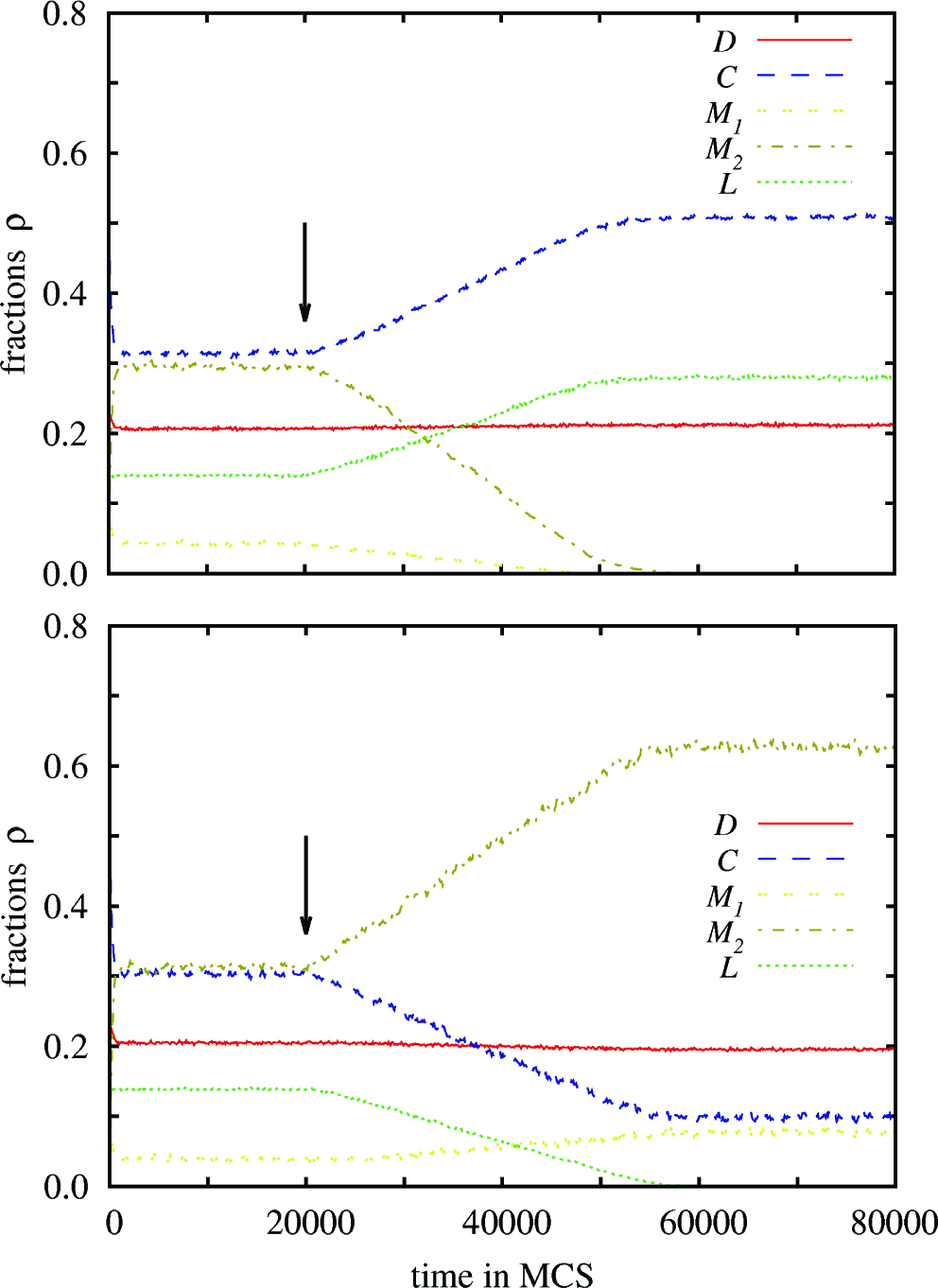
Time evolution of the fraction of strategies, corresponding to the snapshots depicted in Fig. 2. The upper panel uses *r* = 2.80 and *γ* = 0.35, where the winner is the *DCL* phase. The lower panel uses *r* = 2.81 and *γ* = 0.35, where the *DCM*_1_*M*_2_ phase wins. The arrow in both panels at 20000 MCS denotes the time point when the vertical border that separates the two subsystem solutions is removed and the two start competing for space. We have used a 2400 × 2400 square lattice in this case, although the snapshots in Fig. 2 present just a 200 × 200 cutoff of the whole population for clarity (that is also why periodic boundary conditions cannot be inferred there).

## IV. DISCUSSION

To sum up, we have presented the concept of stability of subsystem solutions in agent-based models. We have argued for the necessity of this frequently overlooked approach for the correct and accurate determination of phase transitions, in particular discontinuous phase transitions, in agent-based models where the competing states or strategies each agent can choose from are more than two. Since a subsystem solution can be formed by any subset of all possible agent states, the stability of subsystem solutions is key for attaining certainty that an observed simulation outcome of an agent-based model is actually stable and valid in the large system-size limit. The latter is crucial for the correct determination of phase transitions between different system-wide stable solutions, and for the understanding of the underlying microscopic processes that led to these phase transitions. We emphasize that the described methodology is of the utmost important for teachers and students of complex systems research, and as such should find entry into the appropriate physics curricula at the graduate and, where applicable, also at the undergraduate level.

By using the spatial public goods game with diverse tolerance as an example, we have shown that the 8 competing strategies could form as many as 255 unique subsystem solutions, with no less than 32385 unique pairs to compete against each other in a round-robin tournament. While this is of course an impossible task, it turns out that subsystem solutions that are formed by two or more strategies are rarely individually stable, which significantly reduces the effort that is needed to determine the most stable system-wide solutions and the phase transitions that separate them. We have also emphasized that, before the competition between two individually stable subsystem solutions begins, the two should be fully formed in the sense that they acquire their characteristic spatiotemporal structure (for example traveling or target waves, checkerboard patterns, compact clusters, etc.). A sufficiently long relaxation time is therefore required that must take into account the formation time of both competing subsystem solutions – this is important because the formation of different subsystem solutions is in general characterized by different, and sometimes very different, time scales. Once these conditions are met, representative snapshots of the lattice and time courses of strategy fractions reveal the average moving direction of the invasion front that initially separates the two subsystem solutions, and thus help determine the most stable system-wide solution.

Although the present example draws on spatial evolutionary games and should be of interest to contemporary statistical physics research on the subject [48–63], the concept of stability of subsystem solutions in agent-based models goes far beyond disciplinary boundaries. Whether agents are humans, firms, ants, or ecological entities, whenever more than two states compete in a structured population (represented by a lattice or a network), the stability of subsystem solutions is crucial for a correct and relevant analysis. This is in fact a key distinction that separates ‘simulation research’ from statistical physics research concerning agent-based models and complex systems in general. With today’s computers and programming software, practically any agent-based model is easy to simulate, but the acquisition of correct results requires a careful approach that is seldomly used and advocated for. The current research literature is awash with inaccurate simulation results. The root of the problem lies in overlooked system-size effects and the random extinctions that stem from this, and in particular in the false belief that strategies compete against each other only individually rather than also as subsystem solutions. We hope that this paper will help teachers, students, and practitioners in their future simulation attempts, and that it will also help make the resulting theses and research fit for a physics venue. The time is certainly ripe for the leaps of progress in computer technology and programming software to be followed up by more rigorous simulation practices of agent-based models.

## Acknowledgments

This work was supported by the Slovenian Research Agency (Grants J1-7009 and P5-0027).

## References

[1] P. W. Anderson, Science 177, 393 (1972).

[2] M. Gell-Mann, Engineering and Science 51, 2 (1988).

[3] N. Goldenfeld and L. P. Kadanoff, Science 284, 87 (1999).

[4] Y. Holovatch, R. Kenna, and S. Thurner, Eur. J. Phys. 38, 023002 (2017).

[5] K. Binder, Rep. Prog. Phys. 60, 487 (1997).

[6] H. Hinrichsen, Adv. Phys. 49, 815 (2000).

[7] G. Ódor, Rev. Mod. Phys. 76, 663 (2004).

[8] K. Binder and D. K. Hermann, Monte Carlo Simulations in Statistical Physics (Springer, Heidelberg, 1988).

[9] M. E. J. Newman and G. T. Barkema, Monte Carlo Methods in Statistical Physics (Oxford University Press, Oxford, 1999).

[10] J. Marro and R. Dickman, Nonequilibrium Phase Transitions in Lattice Models (Cambridge University Press, Cambridge, U.K 1999).

[11] D. Landau and K. Binder, A Guide to Monte Carlo Simulations in Statistical Physics (Cambridge University Press, Cambridge 2000).

[12] C. Castellano, S. Fortunato, and V. Loreto, Rev. Mod. Phys. 81, 591 (2009).

[13] S. Galam, Sociophysics: A physicist's modeling of psychopolitical phenomena (Springer, New York, 2012).

[14] M. Perc, Phys. Lett. A 380, 2803 (2016).

[15] M. Perc, J. J. Jordan, D. G. Rand, Z. Wang, S. Boccaletti, and A. Szolnoki, Phys. Rep. 687, 1 (2017).

[16] G. Szabó and G. Fath, Phys. Rep. 446, 97 (2007).

[17] C. P. Roca, J. A. Cuesta, and A. Sanchez, Phys. Life Rev. 6, 208 (2009).

[18] M. Perc and A. Szolnoki, BioSystems 99, 109 (2010).

[19] J. M. Pacheco, V. V. Vasconcelos, and F. C. Santos, Phys. Life Rev. 11, 573 (2014).

[20] Z. Wang, S. Kokubo, M. Jusup, and J. Tanimoto, Phys. Life Rev. 14, 1 (2015).

[21] Z. Wang, L. Wang, A. Szolnoki, and M. Perc, Eur. Phys. J. B 88, 124 (2015).

[22] M. R. D’Orsogna and M. Perc, Phys. Life Rev. 12, 1 (2015).

[23] R. Pastor-Satorras, C. Castellano, P. Van Mieghem, and A. Vespignani, Rev. Mod. Phys. 87, 925 (2015).

[24] Z. Wang, C. T. Bauch, S. Bhattacharyya, A. d’Onofrio, P. Manfredi, M. Perc, N. Perra, M. Salathe, and D. Zhao, Phys. Rep. 664, 1 (2016).

[25] R. Albert and A.-L. Barabasi, Rev. Mod. Phys. 74, 47 (2002).

[26] M. E. J. Newman, SIAM Review 45, 167 (2003).

[27] S. Boccaletti, V. Latora, Y. Moreno, M. Chavez, and D. Hwang, Phys. Rep. 424, 175 (2006).

[28] M. Barthelemy, Phys. Rep. 499, 1 (2011).

[29] E. Estrada, The structure of complex networks: Theory and applications (Oxford University Press, Oxford, 2012).

[30] P. Holme and J. Saramaki, Phys. Rep. 519, 97 (2012).

[31] M. Kivela, A. Arenas, M. Barthelemy, J. P. Gleeson, Y. Moreno, and M. A. Porter, J. Complex Netw. 2, 203 (2014).

[32] S. Boccaletti, G. Bianconi, R. Criado, C. del Genio, J. Gómez-Gardenes, M. Romance, I. Sendina-Nadal, Z. Wang, and M. Zanin, Phys. Rep. 544, 1 (2014).

[33] A.-L. Barabasi, Network Science (Cambridge University Press, Cambridge, 2015).

[34] J. M. Epstein, Complexity 4, 41 (1999).

[35] E. Bonabeau, Proc. Natl. Acad. Sci. USA 99, 7280 (2002).

[36] V. Grimm, E. Revilla, U. Berger, F. Jeltsch, W. M. Mooij, S. F. Railsback, H.-H. Thulke, J. Weiner, T. Wiegand, and D. L. DeAngelis, Science 310, 987 (2005).

[37] J. D. Farmer and D. Foley, Nature 460, 685 (2009).

[38] L. Feng, B. Li, B. Podobnik, T. Preis, and H. E. Stanley, Proc. Natl. Acad. Sci. USA 109, 8388 (2012).

[39] R. M. Axelrod, The complexity of cooperation: Agent-based models of competition and collaboration (Princeton University Press, Princeton, 1997).

[40] C. Adami, J. Schossau, and A. Hintze, Phys. Life Rev. 19, 1 (2016).

[41] C. M. Macal and M. J. North, Journal of Simulation 4, 151 (2010).

[42] M. Perc, Eur. J. Phys. 38, 045801 (2017).

[43] M. Perc, J. Gomez-Gardenes, A. Szolnoki, and L. M. Floria and Y. Moreno, J. R. Soc. Interface 10, 20120997 (2013).

[44] A. Szolnoki and X. Chen, Phys. Rev. E 92, 042813 (2015).

[45] A. Szolnoki and M. Perc, New J. Phys. 18, 083021 (2016).

[46] A. Szolnoki, M. Perc, and G. Szabo, Phys. Rev. E 80, 056109 (2009).

[47] A. Szolnoki, M. Mobilia, L.-L. Jiang, B. Szczesny, A. M. Rucklidge, and M. Perc, J. R. Soc. Interface 11, 20140735 (2014).

[48] J. Tanimoto and N. Kishimoto, Phys. Rev. E 91, 042106 (2015).

[49] J. Vukov, L. Varga, B. Allen, M. A. Nowak, and G. Szabo, Phys. Rev. E 92, 012813 (2015).

[50] H.-X. Yang, Z.-X. Wu, Z. Rong, and Y.-C. Lai, Phys. Rev. E 91, 022121 (2015).

[51] A. Hintze and C. Adami, Phys. Biol. 12, 046005 (2015).

[52] Y. Li, X. Liu, J. C. Claussen, and W. Guo, Phys. Rev. E 91, 062802 (2015).

[53] N. Masuda and F. Fu, F1000Prime Reports 7, 27 (2015).

[54] M. A. Javarone and F. Battiston, J. Stat. Mech. 7, 073404 (2016).

[55] M. A. Javarone, A. Antonioni, and F. Caravelli, EPL 114, 38001 (2016).

[56] A. Aleta, S. Meloni, M. Perc, and Y. Moreno, Phys. Rev. E 94, 062315 (2016).

[57] R. Matsuzawa, J. Tanimoto, and E. Fukuda, Phys. Rev. E 94, 022114 (2016).

[58] M. Pereda, Phys. Rev. E 94, 032314 (2016).

[59] M. A. Amaral, L. Wardil, M. Perc, and J. K. da Silva, Phys. Rev. E 93, 042304 (2016).

[60] F. L. Pinheiro, F. C. Santos, and J. M. Pacheco, Phys. Rev. Lett. 116, 128702 (2016).

[61] M. A. Amaral, M. Perc, L. Wardil, A. Szolnoki, E. J. da Silva Junior, and J. K. da Silva, Phys. Rev. E 95, 032307 (2017).

[62] X. Xu, Z. Rong, Z.-X. Wu, T. Zhou, and C. K. Tse, Phys. Rev. E 95, 052302 (2017).

[63] F. Fu and X. Chen, New J. Phys. 19, 071002 (2017).

